# The Paraphyletic Origins of Genetic Resistance to Cabbage Stem Flea Beetle in *Brassica oleracea*

**DOI:** 10.64898/2025.12.18.695260

**Authors:** Mathieu Tiret, Cyril Falentin, Christine Lariagon, Clarisse Blandin, Mateo Boudet, Damien Rollandez, Antoine Lauverney, Quentin Bazerque, Maria Manzanares-Dauleux, Celine Robert, Sebastien Faure, Antoine Gravot

## Abstract

The cabbage stem flea beetle (CSFB) poses a growing threat to winter *Brassica* crops in Europe, yet the genetic basis of resistance remains poorly understood. To clarify the genetic architecture and evolutionary origins of resistance to CSFB adult feeding, we conducted a genome-wide association study (GWAS) by combining high-throughput pool-sequencing and a large-scale non-choice feeding assay on 113 *Brassica oleracea* accessions from wild (or feral) populations and major domesticated morphotypes. We demonstrate that resistance displays moderate heritability with a predominantly polygenic basis, revealing strong phenotypic divergence among morphotypes: *B. oleracea* var. *capitata* was generally susceptible, whereas var. *botrytis* and wild populations showed markedly higher resistance. Despite this polygenic background, we identified a major-effect candidate QTL on chromosome C01 with strong enrichment of resistance alleles in wild populations and susceptible alleles in var. *capitata*. Genome-wide F-statistics and heterozygosity scans revealed a recent selective sweep at this locus in wild lineages. Considering current evidence for the feral origin of contemporary “wild” populations, our results suggest that resistance evolved after domestication and subsequent feralization, independently of resistance in var. *botrytis*. This paraphyletic distribution underlines the critical importance of integrating demographic history into quantitative genetic analyses of domesticated plant systems.

## Introduction

The cabbage stem flea beetle (*Psylliodes chrysocephala* L.; CSFB) is a long-standing specialist herbivore of *Brassica*ceae and related families, reflecting an evolutionary association that long predates modern agriculture (Gikonyo et al., 2024). Host specialization is pervasive within the *Psylliodes* genus, with nearly half of its species feeding primarily on *Brassicaceae* (Gikonyo et al., 2019). Despite this long co-evolutionary history, the CSFB has only recently become a major threat to *Brassica* crops, causing considerable yield losses on a global scale (Zheng et al., 2020). The recent resurgence of CSFB coincides with rising pest pressure (Andert et al., 2021) and fewer options for chemical controls, together highlighting the need to develop sustainable integrated pest management (Ortega-Ramos et al., 2022).

While the CSFB interacts with *Brassica* crops at various stages, the adult feeding phase represents a particularly critical window in its univoltine life cycle (Bonnemaison & Jourdheuil, 1954; Conrad et al., 2021). In early autumn, upon arrival on *Brassica* fields, CSFB females undergo a 10–15-day pre-oviposition maturity-feeding phase, during which they inflict the characteristic “shotgun hole” damage on young leaves (Thioulouse et al., 1984; Mathiasen et al., 2015; Ortega-Ramos et al., 2022; Tixeront et al., 2024). Although top-down control approaches – such as landscape management or natural predation – contribute to regulation, they have proven insufficient on their own, renewing interest in bottom-up approaches, particularly breeding for genetic resistance to CSFB damage (Pigot, 2023).

Genetic resistance to insect pests, however, presents a persistent challenge for crop improvement. Its genetic architecture is often highly polygenic (Kliebenstein, 2014; Ochoa-Lopez et al., 2018), which disperses effects across many loci and causes linkage disequilibrium (LD) patterns around markers to conflate the causal signal with the population stratification. Without explicit statistical correction and careful sampling, population stratification will in fact lead to spurious association and failure to identify causal variants (Wellenreuther & Hansson, 2016; Tibbs-Cortes et al., 2021). Uncovering the genetic basis of the resistance to CSFB thus requires a deep understanding of the evolutionary history, which is even more challenging in *Brassica* crops, where domestication, feralization, and recent agricultural homogenization adds a complex anthropogenic layer onto natural demographic processes (Mabry et al., 2021).

The lack of strong genetic resistance to adult CSFB feeding documented so far in oilseed rape (*Brassica napus* L.), the most widely cultivated *Brassica* crop in Europe, may be explained by the erosion of resistance during crop domestication, as a result of genetic bottlenecks, selection for agronomic traits, and the relaxation of defense pressures (Turcotte et al., 2014; Chen et al., 2015). Genetic diversity in oilseed rape is in fact limited (Allender & King, 2010), partly due to its recent allopolyploid origin from hybridization between *Brassica oleracea* L. and *Brassica rapa* L. (Chalhoub et al., 2014). In contrast, wild and cultivated cabbages (*B. oleracea*) exhibit extensive morphological and genomic diversity across wild, feral, and domesticated populations (Cheng et al., 2016; Cai et al., 2022) – positioning *B. oleracea* as a potentially rich reservoir of adaptive alleles. Fully exploiting this diversity, however, and identifying the causal variants underlying resistance, requires a clear understanding of its evolutionary history in relation to domestication.

In this study, we hypothesized that *B. oleracea* harbors structured genetic variation underlying resistance to CSFB adult feeding, and that wild populations may retain adaptive alleles that have been diminished or lost in domesticated forms. To test these hypotheses, we conducted a large-scale non-choice feeding experiment on 113 *B. oleracea* accessions from wild populations and major domesticated morphotypes. Most accessions were characterized through high-throughput pool-sequencing (30 plants per accession; 2,926 individuals in total), enabling a robust genome-wide association study (GWAS) of resistance. This design provided a comprehensive view of the genetic architecture of resistance, and revealed that the resistance observed in wild populations cannot be explained solely by a domestication-loss scenario, reflecting instead a more complex evolutionary trajectory.

## Materials and Methods

### Plant Material

A total of 113 *B. oleracea* accessions were collected, comprising both cultivated landraces and wild populations. The collection includes accessions belonging to: *B. oleracea* var. *acephala* (kale), var. *botrytis* (cauliflower), var. *capitata* (cabbage), var. *italica* (broccoli), accessions of an unknown morphotype, and wild populations (rather feral; Mabry et al., 2021; Cai et al., 2022) hereafter referred to as var. *oleracea* (Table 1). The collection comprises 90 accessions that originated from the BrasExplor project (Falentin et al., 2024), 7 from the worldwide core collection assembled within the PROBIODIV project (Falentin et al., 2025), one reference accession (*B. oleracea* var. *italica* ‘HDEM’), 9 accessions from the Wageningen WUR genebank (‘CGN06903’, ‘CGN07077’, ‘CGN07123’, ‘CGN11073’, ‘CGN11132’, ‘CGN14063’, ‘CGN15129’, ‘CGN18431’, ‘CGN18947’), and 6 accessions from the Warwick HRI genebank (‘HRIGRU 6431’, ‘HRIGRU 7343’, ‘HRIGRU 7795’, ‘HRIGRU 8658’, ‘HRIGRU 8694’, ‘HRIGRU 9846’) (details in Table S1). Accessions were selected from public germplasm collections and breeding resources to maximize phylogenetic and morphological diversity while maintaining multiple representatives within each morphotype. This sampling strategy aimed to capture the principal domestication lineages of *B. oleracea* and to enable comparisons between cultivated and wild gene pools.

**Table 1.**
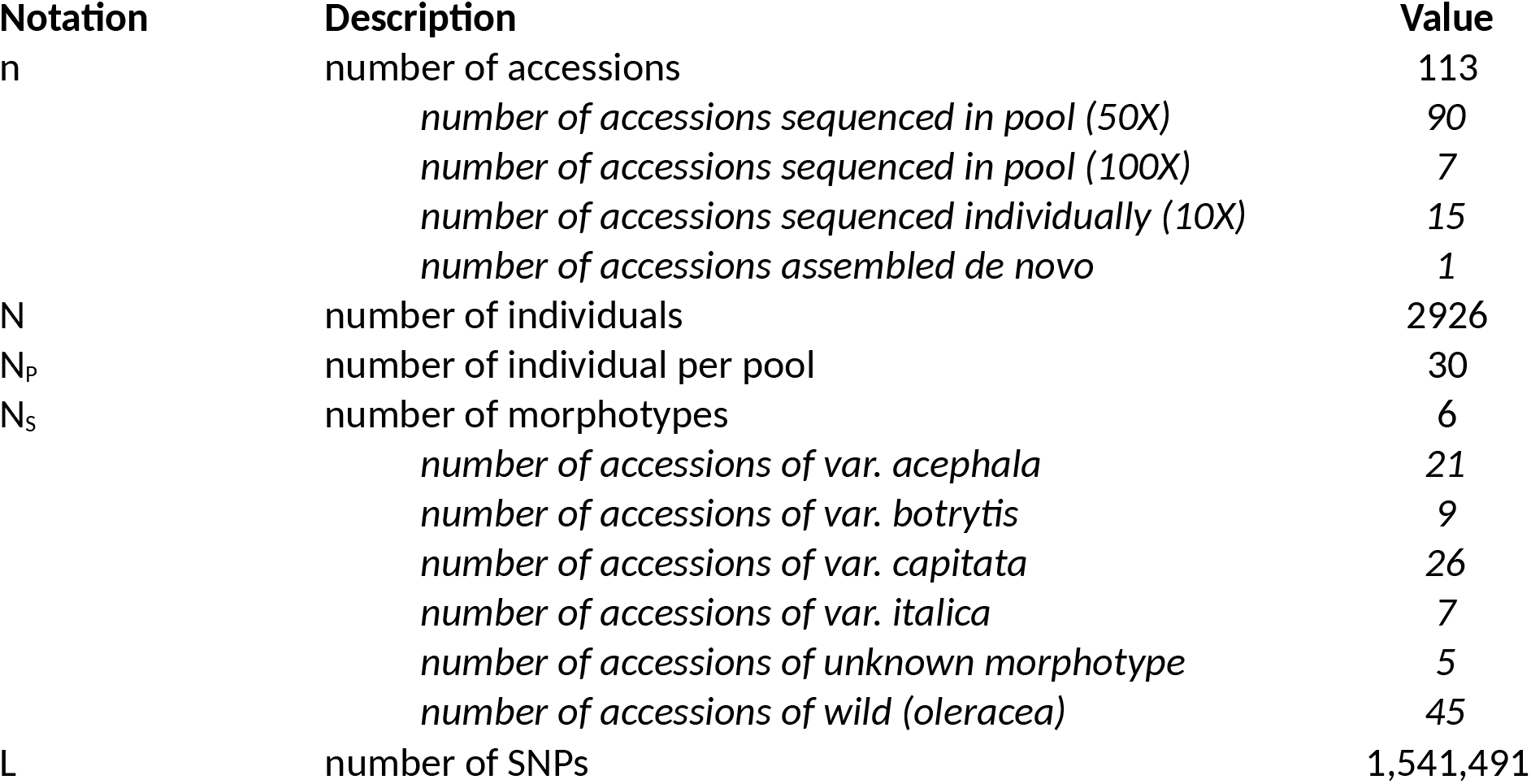
detailed figures of the experiment.

Seed lots used for phenotyping were, when possible, regenerated through controlled multiplication campaigns conducted at Ploudaniel (France; 48°30’06.5”N 4°19’33.6”W). To best convey the within-accession diversity, the largest feasible number of parental plants was used in each multiplication. For the BrasExplor accessions, seeds were obtained in 2021 from pairwise crosses among six parental plants per accession. The seven PROBIODIV accessions were multiplied in 2023 using 2 plants each. The ‘HDEM’ cultivar line was regenerated through single-seed descent in 2022. For the fifteen genebank accessions, original seed lots supplied by the repositories were used directly without further multiplication.

### Genetic data

Whole-genome resequencing data were obtained for all accessions using seed samples collected prior to multiplication, in order to mimic an F_2_-like design. For the BrasExplor panel, DNA was extracted from pools (i.e., pool-seq) of 30 individuals per accession following the protocols described in Maillet et al. (2023) and Tiret et al. (2024). Libraries were sequenced using Illumina short-read paired-end technology (150 bp; Illumina Inc., San Diego, CA, USA) with a target coverage of 50× (details in Tiret et al., 2024). Accessions from the PROBIODIV project were processed with the same protocol and pooled-sequenced (30 individuals per accession) at a target coverage of 100×. Genebank accessions were resequenced individually (single-seed DNA) using the same platform and read length, with a target coverage of 10×. For the ‘HDEM’ reference genome, resequencing data were derived from its assembled genome sequence. An additional accession of *Brassica montana* L. (‘BM_F_SOUL_W_A’; Falentin et al., 2024), a close relative of B. oleracea sharing the same karyotype (Arias & Pires, 2012; Saban et al., 2023), was included as an outgroup to root the phylogenetic tree. In total, the sequencing dataset represents 2,926 individuals (97 pools of 30 individuals each plus 16 single-seed resequenced accessions).

Raw reads (FASTQ files) were processed using fastp v1.0.1 (Chen, 2025) for quality control. The pipeline included trimming of poly-G and poly-X tails, removal of reads shorter than 30 bp, trimming of bases with Phred quality below Q20 at both ends, and removal of Illumina TruSeq adapter sequences according to the manufacturer’s recommendations. Clean reads were mapped to the ‘HDEM’ reference genome (Belser et al., 2018) using BWA-MEM2 v2.3 (Vasimuddin et al., 2019) with the parameter settings recommended by Regier et al. (2018). Duplicate reads were removed using SAMtools v1.22.1 (Danecek et al., 2021). Variant calling was performed with BCFtools v1.22 using the ‘-X illumina’ option and a maximum read depth of 1000 (‘-d 1000’). Variants were filtered with BCFtools to retain only high-quality biallelic SNPs and INDELs, with more than 5 accessions at both extreme frequencies (below 0.1 and above 0.9) to avoid leverage effects in GWAS, and a missing rate < 5% (details in Supplementary Materials S1). The centromeric regions of chromosomes C01 and C9 displayed abnormally high sequencing coverage; consequently, SNPs located on C01 between 26-31.5 Mb and on C9 between 30.5-38 Mb were excluded from all analyses (Fig. S1). The final dataset comprised 1,541,491 SNPs. The VCF file was then converted into a CSV file using PLINK2 v2.0.0 (Chang et al., 2015), where pool individuals were represented with their estimated allele frequencies (VAF field), and 0/1/2 for individuals (the genebank accessions) – all VAFs were multiplied by 2 to match the scale, and was modulated by step of 1/30 (to represent the 30 individuals/pool). This matrix [L x n], denoted X, will be referred to as the pool genotype table. All subsequent statistical analyses were implemented in R v4.3.3 (R Core Team, 2024).

### Population structure

Alleles were polarized using the accession of *B. montana* as an outgroup, replacing the reference allele of ‘HDEM’ with that of the outgroup wherever they differed. This provided an ancestral/derived orientation for all variants across the *B. oleracea* accessions. At each SNP, the allele frequency was estimated as the weighted mean accounting for the number of individuals per pool (1 or 30). Genetic relationships among accessions were visualized using Principal Coordinates Analysis (PCoA). Consistent with previous work showing that *B. oleracea* population structure aligns with morphotype differentiation (from passport data), accessions were assigned to six pre-defined clusters: *acephala, botrytis, capitata, italica, oleracea* (wild), and unknown.

To capture local genomic signals of differentiation, hierarchical F-statistics, mainly F_GT_, which is an equivalent of F_ST_ for between-cluster divergence when dealing with pools (Gautier et al., 2024), were estimated along the genome using sliding windows of 35 SNPs with the ‘poolfstat’ R package (v3.0.0; Gautier et al., 2024). We additionally computed the D_i_ statistic per genetic cluster by computing the F_GT_ between the focal cluster and the rest, and scaling the result (Akey et al., 2010; McQuillan et al., 2022). Finally, analogous F-statistics were computed for phenotypic groups: resistant (R, first quartile), intermediate (I, second and third quartiles), and susceptible (S, fourth quartile); this distinction, hereafter denoted R/S, allowed us to quantify genetic differentiation between resistant and susceptible accessions.

### Genomic Relationship matrix

The additive genomic relationship matrix (G) was computed from the pool genotype table (X) following the first method of VanRaden (2008). Briefly, allele counts in X were centered by their estimated allele frequencies and scaled by the corresponding heterozygosity, ensuring unit variance across loci. To represent genetic relatedness among morphotypes, we also derived a Q matrix by applying the same method to an aggregated genotype matrix (X^S^), obtained by averaging the rows of X within each morphotype. The matrix Q was rescaled to have a unit mean on the diagonal to match G, and for simplicity, is still referred to as Q.

The dominance relationship matrix (D) was parameterized following Vitezica et al. (2013). For each locus i and individual (or pool) j, we defined the dominance-coded value as: - X_ij_^2^ + (1+2p_i_).X_ij_ – 2p_i_ ^2^, where X_ij_ is the value of the i-th locus for the j-th individual (0-2 scale), and p_i_ the estimated MAF at the i-th locus. This formulation ensures that additive and dominance components are orthogonal under Hardy–Weinberg equilibrium. Because empirical correlations between G and D persisted in the pooled dataset, we enforced orthogonality through projection (as in Vitezica et al., 2013): Do = D – G.tr(DG)/tr(GG), where *tr* denotes the matrix trace. The resulting matrix Do was also rescaled to have a unit mean on the diagonal, and is hereafter referred to as D. Since the present dataset is based on pool-seq allele frequencies, the dominance term should be interpreted as reflecting shared excess heterozygosity across accessions rather than individual-level dominance deviations – the objective here is to estimate the dominance variance.

### Phylogeny

Distance-based phylogenetic trees were constructed to examine how resistance segregated across the evolutionary path of each accession. The tree was inferred from the G matrix after converting it to pairwise distances using sqrt(Gii + Gjj – 2Gij), as this measure yielded phylogenetic relationships most consistent with previous findings (Cai et al., 2022; see Supplementary Materials S2). Phylogeny reconstruction was performed with the FASTME algorithm (ordinary least square), using the ‘ape’ R package (v5.8.1; Paradis & Schliep, 2019). Nodes with branch lengths below 0.001 were collapsed, and the resulting tree was rooted to the *B. montana* outgroup. The segregation of resistance was projected onto the phylogeny using ancestral state reconstruction, using the ‘phytools’ R package (v2.5.2; Revell, 2024), following the approach described by Bravo et al. (2019). The annotated tree was visualized with the ‘ggplot2’ (v4.0.0; Wickham, 2016) and ‘ggtree’ (v3.99.2; Yu, 2022) R packages.

### Experimental design & Phenotype data

The phenotype was defined as the leaf area consumed by adult CSFB under a non-choice feeding assay (‘shotgun holes’ leaf damage). This metric provides a quantitative estimate of plant resistance to feeding but does not distinguish among potential defense mechanisms such as antibiosis, antixenosis, or tolerance. The *B. napus* ‘Aviso’ and *B. rapa* ‘Malwira’ were used as the susceptible controls (Döring & Ulber, 2020; Li et al., 2024), and *Sinapis alba* L. ‘Sarah’ as the resistant control. Diapaused adult CSFB (minimum 90 days old), reared on-site under controlled conditions (12h/12h photoperiod, 16°C/10°C), were used for all feeding experiment. The CSFB collection was supplemented with field samples collected from Le Rheu (France) in fall 2024 combined with individuals from a rearing colony maintained in Mondonville (France, IN-NOLEA) that is regularly supplemented with local field samples to maintain genetic diversity. After a 72-hour fasting period, two (unsexed) beetles were placed individually in tubes containing a single plant at the 2–3 leaf stage (ca. three weeks old, i.e., the typical stage at which CSFB causes damage under field condition) grown in mini-pots (Fig. S2). Cotyledons were removed 72 hours before exposure to standardize the feeding substrate and restrict insect feeding to the first true leaf.. The experiment lasted 48 hours in a climate chamber (16h/8h photoperiod, 21°C/18°C). Leaves were then excised, flattened, and photographed on a backlit surface. Image analysis was performed with a semi-automated script in FIJI v2.9.0 (Schindelin et al., 2012) requiring manual delineation of external damaged areas to quantify the leaf area consumed (mm^2^). This setup enabled the simultaneous screening of all accessions, with each replicate considered as a block. After data curation (removal of 0-damage leaves, outliers, plants with dead beetle during the experiment, or unexploitable picture), the final dataset included 742 observations across 113 genotypes (6.566 ± 1.085 replicates per genotype) within nine exploitable blocks, corresponding to a fill rate of 72.9% (see Supplementary Materials S3). The data was treated as obtained from an incomplete randomized block design due to missing data. To reduce positive skewness, the phenotype was square-root transformed. For consistency, the transformed trait is hereafter expressed in mm and referred to as ‘leaf damage’.

### Genomic BLUP

Breeding values of the accessions were estimated using a Genomic Best Linear Unbiased Predictor (GBLUP) under a QK model defined as:

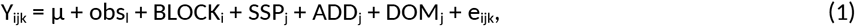

where ‘Y_ijk_’ is the phenotype of the k-th replicate of the j-th accession in the i-th block observed by the l-th observer, μ is the grand mean, ‘obs_l_’ the fixed effect of the l-th observer, ‘BLOCK_i_’ the random effect of the i-th block, ‘SSP_j_’ the random effect associated with the genetic cluster of the j-th accession, ‘ADD_j_’ the random effect of the polygenic additive effect, ‘DOM_j_’ the random effect of the dominance deviation, and ‘e_ijk_’ the residual (see details in Supplementary Materials S4). Both the block effect and the residuals follow a Gaussian distribution with diagonal covariance matrices, and the ‘SSP’, ‘ADD’ and ‘DOM’ effects follow a Gaussian distribution with a covariance matrix equal to σ_Q_^2^Q, σ_G_^2^G, and σ_D_^2^D respectively. All random effects and residuals were assumed independent. Model (1) was fitted using the ‘mmes()’ function (R package ‘sommer’ v.4.4.3; Covarrubias-Pazaran, 2016), which uses a REML algorithm to fit the variance components; we additionally provided a relaxed tolerance parameter for the matrix inversion (tolParInv = 0.5) to facilitate convergence.

Because additive and dominance effects exhibited partial collinearity, we implemented a two-step estimation procedure to obtain stable and interpretable additive effects. In a first step, model (1) was used to estimate dominance effects, which were then subtracted from the raw phenotype (Y* = Y – DOM). The adjusted phenotype was subsequently analyzed with a reduced QK model:

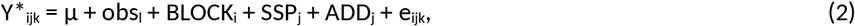

with indices and factors defined as above. Model (2) was fitted using the ‘mmes()’ function under default settings. This two-step approach limited confounding between additive and dominance components and provided a more conservative estimation of additive genetic values. Prediction accuracy of each accession was computed as r_j_ = 1 – PEV_j_/σ_G_^2^G_jj_, following Misztal et al. (2002) and the implementation of BLUPF90, where PEV_j_ is the prediction error variance of the j-th accession and G_jj_ the j-th diagonal element of the genomic relationship matrix. Population differentiation in additive effects was quantified as 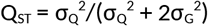 following Spitze (1993), and compared to the empirical distribution of neutral differentiation values as recommended by Whitlock (2008); we will use the F_GT_ measure instead of F_ST_ to account for hierarchical clustering (Gautier et al., 2024).

Model robustness was evaluated by a bootstrap resampling of each observation (1,000 replicates), also limiting the GBLUP shrinkage. For each estimate, we summarized the bootstrap mean, variance, skewness (R package ‘e1071’ v1.7.16; Meyer et al., 2024) and 95% confidence intervals. Robustness was quantified as the absolute skewness across bootstrap replicates, with values <1 considered robust and <2 moderately robust. Because the relaxed tolerance required to invert large covariance matrices may induce occasional numerical instability, the entire two-step estimation pipeline was repeated until all 1,000 bootstrap replicates satisfied a convergence criterion of <30% deviation in the estimated genetic variance between model (1) and model (2). Permutation tests were conducted by shuffling accessions within each morphotype – thereby preserving both the phenotype-environment association and the genotype-morphotype structure – and repeated 1,000 times. To compare the bootstrap and permutation distributions, we quantified their separation using Cohen’s D effect size (function ‘cohen.d()’, R package ‘effsize’ v0.8.1; Torchiano, 2020). This metric was preferred over the Kolmogorov–Smirnov or chi-square tests, which become overly susceptible in the presence of large sample sizes (1,000 replicates).

### GENOME-Wide Association Study

The pool-seq GWAS was conducted by back-solving SNP effects from GBLUPs following the mixed-model equations of Aguilar et al. (2019), implemented in a custom R script (see Supplementary Materials S5). For each SNP, we computed the allelic effect, its standard error, and the associated p-value. SNP-specific coefficients of determination (R^2^) were computed similarly than in the ‘TwoSampleMR’ R package (v0.6.24; Hemani et al., 2018). Enrichment of certain genomic region in “resistant” alleles was tested using a weighted chi-squared test, where each SNP was weighted by its R^2^ value, using the ‘weights’ R package (v1.1.2; Pasek, 2025).

As in the GBLUP analyses, SNP-effect estimation was bootstrapped (1,000 replicates). Confidence intervals for genome-wide quantities (allelic effects, R^2^, p-values) were approximated under a Gaussian assumption. The proportion of variance explained by each chromosome was computed as the sum of the chromosome’s SNP R^2^ divided by the overall sum of R^2^. Significance thresholds for GWAS were determined using the ‘maxT’ permutation framework (John et al., 2024), yielding a 95% threshold of 6.724, compared with the more conservative Bonferroni value of 7.489. Genomic inflation factors (λ_median_) was computed as in Listgarten et al. (2012).

To evaluate the stability of allelic effects, we carried out an independent validation experiment on six genotypes selected for highly contrasted resistance levels (see Supplementary Materials S6). Predictive ability (PA) was quantified as the Pearson correlation between phenotypes and genomic predictions computed from the estimated additive and dominance effects: PA = ρ(Y_2_, Xa + d + s), where ‘Y_2’_ is the observed phenotype ‘a’ the estimated allelic effects, ‘d’ the estimated dominance deviation, ‘s’ the estimated morphotype effect, and X the pool genotype table. Similarly, we conducted a leave-one-morphotype-out genomic prediction, whereby the model was fit on all accessions but those of one morphotype, and stability was assessed via the PA on the withheld morpho - type accessions.

Based on the GWAS results, we defined three SNP subsets: (i) the full genome-wide SNP set; (ii) a reduced panel of 200,000 SNPs with the largest effects; (iii) a QTL-based SNP set defined as ±0.2 Mb around high-score peaks. To investigate the relationship between demographic history and the genetic architecture of resistance, we computed for each SNP subset a weighted genomic relationship matrix (GRM) following the first method of VanRaden (2008), scaling the numerator by SNP-specific R^2^ values.

## Results

Leaf damage exhibited considerable variation among *B. oleracea* accessions, averaging 4.092 ± 1.950 mm (Fig. 1a). The damage observed in *B. oleracea* fell below the levels of both *B. napus* (4.442 ± 2.211 mm) and *B. rapa* (6.409 ± 2.081 mm), yet exceeded that of *S. alba* (2.055 ± 1.326 mm). Within *B. oleracea*, the most resistant landrace, ‘BO_I_SOLE_L_C’, recorded a leaf damage of 2.829 ± 0.349 mm, closely followed by the most resistant wild population, ‘BO_E_LARE_W_A’, at 2.841 ± 0.297 mm. In contrast, the least resistant landrace, ‘BO_F_DOUA_L_B’, reached a leaf damage of 5.457 ± 0.349 mm.

**Figure 1.**
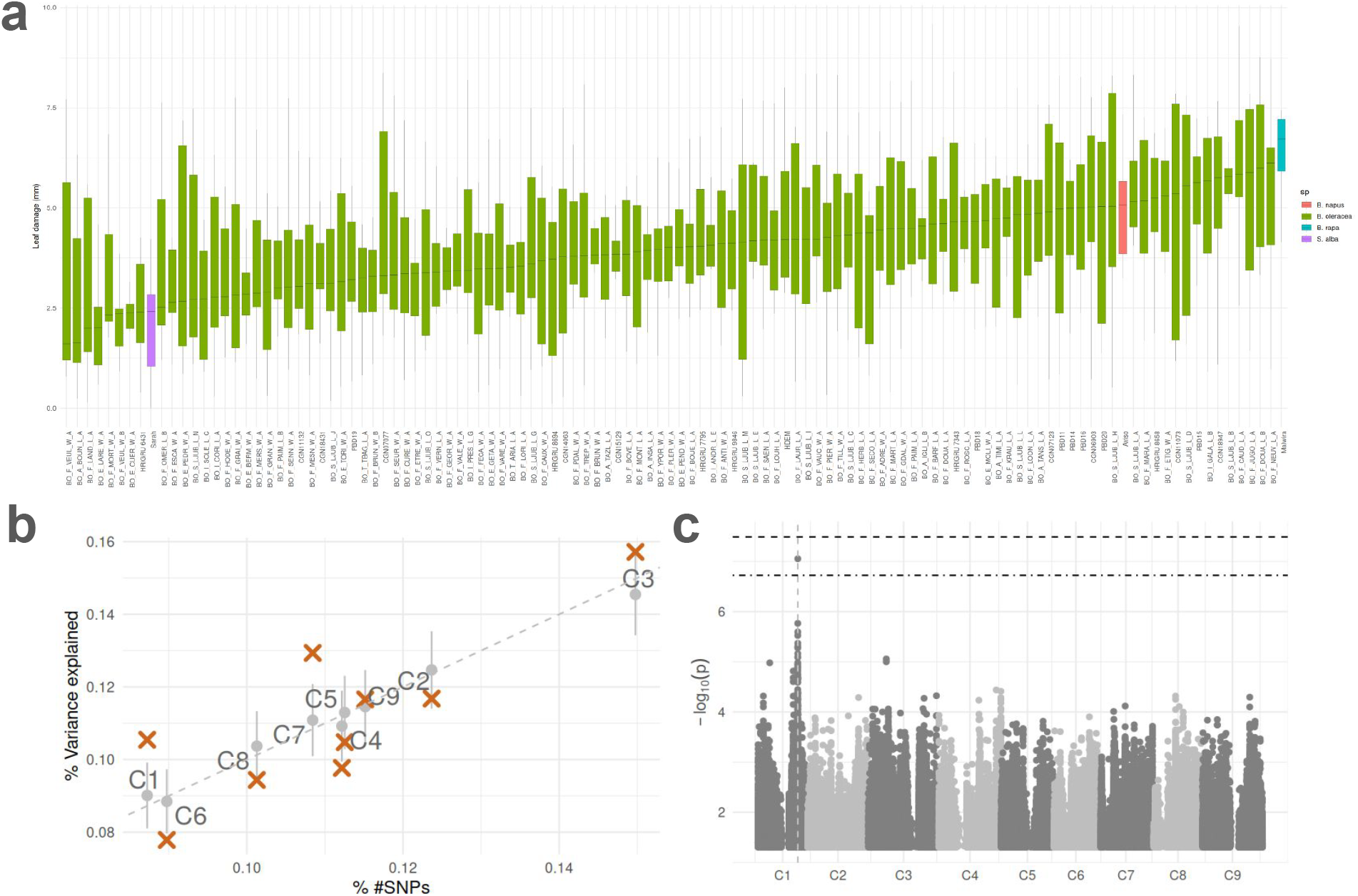
Genetic architecture of the resistance to CSFB in *B. oleracea*. **(a)** Distribution of the leaf damage (mm) across accessions of *B. oleracea* (green), *B. napus* (red), *B. rapa* (blue), and *S. alba* (purple). **(b)** Percentage of variance explained per chromosome, with the full SNP set (grey circles) or the top 200,000 SNPs (red crosses). **(c)** Manhattan plot of the GWAS, with the permutation threshold (alternate dotted line) and the Bonferroni threshold (dotted line).

### 1. Polygenic architecture of resistance to leaf damage

After accounting for dominance – which explained 21.9% of phenotypic variation – the variance decomposition indicated that unexplained environmental effects were the major source of variation (∼80%; Table 2). Polygenic effects contributed 15.7%, and morphotype effects 3.5%. The overall model fit remained modest (R^2^ = 0.328), consistent with strong within-plot heterogeneity likely reflecting the behavioral and stochastic nature of CSFB feeding. Despite this noise, heritability was significantly positive (h^2^ = 0.157), confirming an additive genetic component underlying this trait. GBLUP coefficient of determination (CD) values were generally high (mean 0.675 [0.501; 0.798]), confirming reliable discrimination among accessions. Residual diagnostics showed no systematic biases, and bootstrap analyses indicated stable variance estimates (|skew| < 0.819), as well as for GBLUPs (|skew| < 0.374). On low-CD genotypes, slight left-tail skews were estimated, suggesting that outlier observations were events of unexpectedly low consumption.

**Table 2.** Coefficient of determination, variance and morphotype effects: estimated values, percentage of explained variance, 95% confidence intervals, and robustness score across bootstraps. All effects were estimated with model (2), after correction of the dominance effect (21.9%).

|  | Estimated value | % | 95% CI | \|skew\| |
| --- | --- | --- | --- | --- |
| $R^2$ | 0.328 | | [0.284; 0.370] | 0.059 |
| *Variance* |  |  |  |  |
| $\sigma_G^2$ | 0.635 | 15.7% | [0.454; 0.845] | 0.475 |
| $\sigma_Q^2$ | 0.143 | 3.5% | [0.001; 0.380] | 0.819 |
| $\sigma^2(\text{BLOCK})$ | 0.706 | 17.5% | [0.568; 0.864] | 0.227 |
| $\sigma^2(\text{Residual})$ | 2.555 | 63.3% | [2.382; 2.722] | 0.003 |
| *Random effects* |  |  |  |  |
| *acephala* | 0.130 |  | [-0.009; 0.394] | 0.840 |
| *botrytis* | -0.196 |  | [-0.660; 0.005] | 1.306 |
| *capitata* | 0.272 |  | [0.004; 0.759] | 1.010 |
| *italica* | -0.027 |  | [-0.276; 0.141] | 1.173 |
| *oleracea* | -0.172 |  | [-0.372; -0.003] | 0.244 |
| *unknown* | -0.007 |  | [-0.157; 0.101] | 1.019 |

Allelic effects were globally weak, with SNP-wise R^2^ values averaging 0.009 [0.002; 0.034], consistent with a highly polygenic trait. Chromosome-wide contributions matched SNP density, except for C01, C03 and C05, which displayed enrichment above infinitesimal expectations (Fig. 1b). After correcting for population structure – which slightly over-corrected the test statistics (λ_median_ = 0.747; Fig. S3) – the GWAS identified a single significant SNP on C01 at 41,813,754 bp (an INDEL T/TA), with a significance of −log_10_(p) = 7.054, surpassing the empirical permutation threshold of 6.724 (Fig. 1c). This candidate QTL explained moderate variation (R^2^ = 0.173 [0.115; 0.230]) and displayed robust bootstrap behavior (|skew| = 0.230). Without dominance correction, however, the candidate QTL fell below the significance threshold, indicating that dominance-related genome-wide structure obscured additive signals. The two-step approach employed here ensures that the significant signal conveys an additive effect.

To further assess robustness, genomic prediction was tested on six contrasted genotypes in an independent experiment. Prediction accuracy was high (PA = 0.732 [0.465; 0.904]) with a large effect size compared to permutation (Cohen’s D = 1.829; Fig. S4), confirming that estimated SNP effects capture meaningful signals. When using only the top 200k SNPs, PA was still significantly higher than random (D = 1.572) but reduced to 0.687 [0.464; 0.853], and progressively smaller SNP sets led to drastic decrease in predictive abilities (Fig. S5), reinforcing the hypothesis of a high polygenicity of the trait. For the C01 QTL, despite its statistical significance, by itself had a low predictive power (PA= 0.474 [0.173; 0.757]) and a small effect size (D = 0.504).

### 2. Quantitative susceptibility recently evolved in var. *capitata*

Population structure is an inherent concern for polygenic traits, and in *B. oleracea* the structure aligned strongly with morphotypes (Fig. 2a). Although morphotypes explained only a modest fraction of the phenotypic variance (3.5%), its effect was statistically significant. Within this component, var. *botrytis* and var. *oleracea* tended to show higher resistance (significantly for the latter), while var. *capitata* was the only morphotype displaying a significantly high susceptibility (Table 2). The robustness of these morphotype estimates was moderate (|skew| > 0.840 for most morphotypes), with the exception of var. *oleracea* with high robustness (|skew| = 0.244).

**Figure 2.**
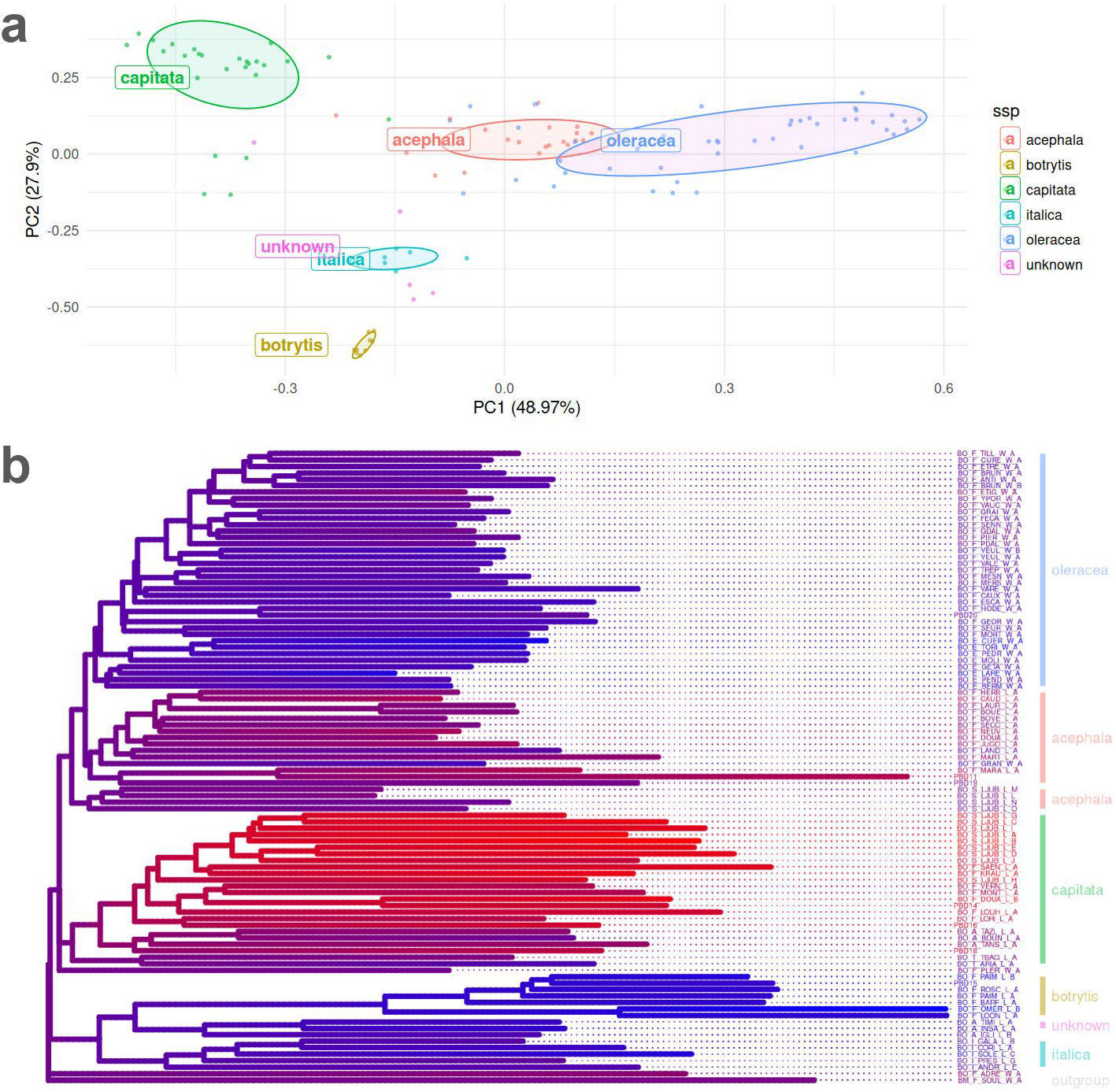
Phylogeny of the resistance to CSFB. **(a)** PCoA of the genetic clustering per morphotype: *acephala* (red), *botrytis* (orange), *capitata* (green), *italica* (cyan), *oleracea* (blue), and unknown (pink). **(b)** GBLUP projected on the phylogenetic tree, with high resistance (blue) and low resistance (red).

When populations are genetically structured, GWAS signals can partly reflect spurious associations rather than true causal variants. To evaluate how robust our inference framework was to this issue, we carried out leave-one-morphotype-out genomic predictions. Predictive ability dropped only marginally (<0.02) when relying on the largest-effect SNP subset, but sharply (>0.26) when using the full SNP set. This contrast shows that the largest-effect SNPs most likely capture the genetic basis of the phenotype that is common across morphotypes, whereas the remaining SNPs capture a more morphotype-specific signal – or reflects overfitting. In other words, the largest-effect SNP set is the part of the resistance signal that is conserved across morphotypes, where the rest is population-specific, does not generalize beyond its own lineage, but enhances PA. This genetic architecture with two layers, one that is shared across morphotypes and another that is specific within each morphotype, is consistent with the Q_ST_–F_GT_ comparison: quantitative divergence for the trait (Q_ST_ = 0.098 [0.001; 0.223]) did not exceed neutral differentiation (F_GT_ = 0.145 [0.002; 0.453]), and the effect size between the two distributions was small (D = 0.416; Fig. S6).

Projecting genomic prediction of resistance (derived from the largest-effect SNP set) onto the inferred phylogenetic tree clarified its evolutionary trajectory (Fig. 2b): the level of resistance is mainly shared within a morphotype, but those who are resistant are not necessarily closely linked, i.e., a paraphyletic resistance. High susceptibility constitutes a derived condition largely confined to var. *capitata*, even though some accessions from other morphotypes (e.g., var. *acephala*) were also susceptible (Fig. 2b). Reconstructed ancestral lineages (older branches close to the root) exhibited only moderate resistance (Fig. 2b), and wild accessions were not the most resistant – counter to expectations under a scenario of domestication-driven erosion of defense. The major C01 QTL followed the same distribution across morphotypes (Fig. S7). Taken together, the segregation of both the QTL and the largest-effect SNP set shows that resistance-associated variants’ evolutionary dynamics did not align with the phylogeny or domestication history: landraces are not uniformly more susceptible than wild accessions.

### 3. Positive selection on resistance-linked alleles in wild populations

Although domestication did not diminish resistance, the allele frequency patterns around the major QTL revealed a contrast between landraces and wild lineages: the latter carried a significantly higher proportion of resistance-associated alleles at the C01 QTL (and its flanking region) than expected from their genome-wide R/S allele composition (weighted χ^2^, p < 0.001; Fig. 3a). On the opposite, var. *capitata* exhibited a significant enrichment of susceptible alleles (p < 0.001; Fig. 3b). The robustness of this QTL was confirmed with a genome-wide F_GT_ scan (Fig. 3c) – a metric that does not dependent on the two-step dominance correction – thereby reinforcing the detected signal: when accessions were grouped by the level of resistance, a significantly strong peak appeared precisely at the C01 QTL (F_GT_ = 0.143), matching the 95% permutation threshold (0.143). This peak vanished when grouping accessions by morphotype (Fig. S8), showing that the differentiation at this QTL is tied to resistance rather than to overall population structure.

**Figure 3.**
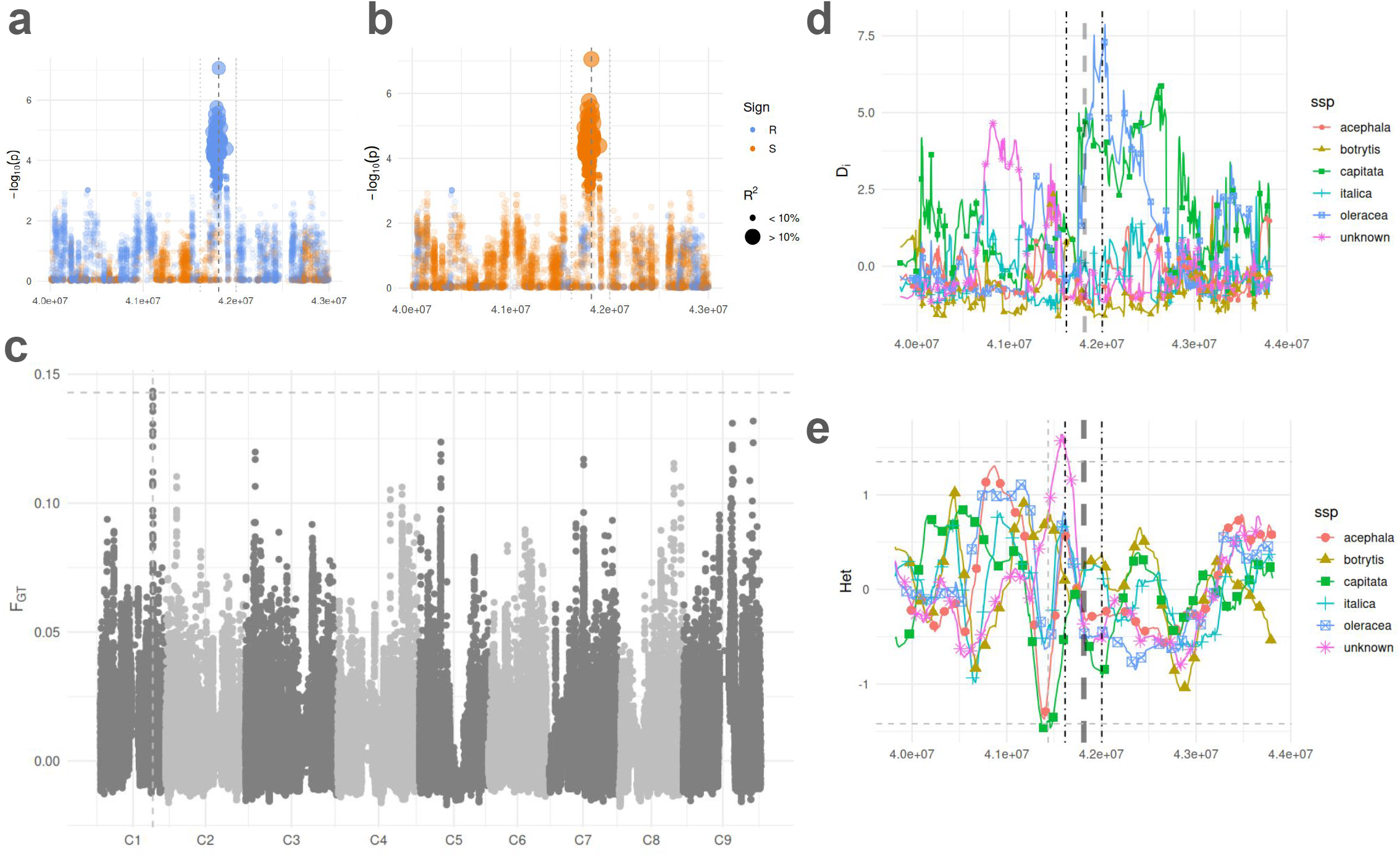
Selective sweep on the C01 QTL. **(a)** Manhattan plot of the GWAS, zoomed into the vicinity of the C01 QTL. Each dot is a SNP, and is polarized according to the average allelic frequencies of var. *oleracea*: orange when the version carried on average by *oleracea* is susceptible, blue otherwise. **(b)** Same as (a), but polarized according to the average frequencies of var. *capitata*. **(c)** FGT scan when clustered by R/S group, showcasing a significant peak on the C01 QTL, surpassing the permutation threshold (dotted line). **(d)** D_i_ scan per morphotype in the vicinity of the C01 QTL: *acephala* (red), *botrytis* (orange), *capitata* (green), *italica* (cyan), *oleracea* (blue), and unknown (pink). **(e)** Same as (d), for heterozygosity scan.

The pairwise F-statistics scan revealed a broad region of elevated D_i_ adjacent to the QTL (42.0– 42.6 Mb), with the strongest contrasts involving var. *oleracea* and var. *capitata* (Fig. 3d). A heterozygosity scan pinpointed that the wild cabbages were the driver of this signal, as they showed a marked reduction in diversity in this genomic region, whereas diversity remained comparatively high in var. *capitata* (Fig. 3e). This combination of high D_i_ and reduced diversity is a signature of positive selection on var. *oleracea*, adjacent to – but not precisely on – the QTL peak.

## Discussion

The present investigation – despite a substantial proportion of unexplained variance in leaf damage, likely reflecting the inherently stochastic aspect of feeding behaviors of CSFB – reveals clear inter-specific contrasts: *B. oleracea* consistently exhibited lower levels of damage than *B. napus* and *B. rapa*, although still more than *S. alba*, a species widely recognized for robust defenses (Hennies, 2016; Döring, & Ulber, 2020). Within *B. oleracea*, genome-wide analyses indicate a moderately heritable trait (h^2^ = 0.157) shaped by a highly polygenic architecture. Yet, embedded in this diffuse background, we identified a major-effect locus on chromosome C01 (∼41.8 Mb) exhibiting a strong signal of recent selection in wild populations. This pattern points toward a dynamic interplay between domestication and diversification in structuring the contemporary distribution of resistance alleles, suggesting that resistance has not been shaped by a simple loss in domesticated lines but rather by successive shifts depending on the cultural and ecological environment.

### 1. Post-feralization selection likely enhanced resistance in wild populations

The feral origin of many European “wild” *B. oleracea* populations (Mabry et al. 2021; Cai et al., 2022) is a critical point to understand the evolutionary trajectory of the resistance to CSFB. Our results indicate that the C01 candidate QTL is significantly associated with the high resistance observed in feral lineages, even though these lineages likely diverged only recently from landraces – most plausibly from var. *acephala* or var. *capitata* (Cai et al., 2022; Saban et al., 2023). The pattern suggests that ancestral populations were moderately resistant at best, and that the variance observed today emerged from divergent pressures acting after domestication: feral lineages experienced renewed selection favoring defense traits (e.g., chemical defense; Poelman et al., 2009), where CSFB pressure no more encountered trade-offs with economically interesting traits. On the opposite, the sweep-like signature on C01 in feral lineages – but not in domesticated morphotypes – is consistent with the hypothesis that domesticated forms experienced relaxed selection on defense likely due to agricultural buffering (e.g., weed control, fertilization, reduced herbivore pressure). This trajectory implies that resistance is not a conserved ancestral trait that was simply lost during domestication but in fact a trait that was actively re-assembled in natural and semi-natural contexts (Kareiva et al., 2007).

The C01 signal lies near a domestication-associated peak reported by Saban et al. (2023; ∼45.2 Mb), but its behavior in our data depended strongly on how populations were grouped. When grouped by morphotype, the signal collapsed; when grouped by resistance phenotype, the signal reappeared. This implies that the resistance, rather than genetic clustering, drives the evolution of this region – but it is indeed tightly linked to the domestication history, and this genomic region requires further investigation. Although our pool-seq data are insufficient for haplotype reconstruction, the wide D_i_ peak is compatible with a recent sweep; if tied to feralization date, its origin should not predate ∼3.5 kya, the estimated divergence between var. *capitata*/*acephala* and var. *botrytis*/*italica* (Saban et al. 2023). Full resolution of the sweep’s age, its role in domestication, and structure will require targeted individual-level resequencing.

The susceptibility of var. *capitata*, which appears to accumulate genome-wide liabilities to CSFB, might be the result of different domestication-driven trade-offs. A plausible scenario involves trade-offs with defense pathways, particularly those linked to glucosinolate biosynthesis and hydrolysis. Breeding for reduced bitterness and improved palatability can inadvertently suppress alleles that contribute to herbivore resistance (Koritsas et al. 1991; Beran et al. 2018). Yet, the relationship between glucosinolates and CSFB is complex: no single compound or class has yet been reported to consistently predict resistance (Bartlet et al., 1996; Döring & Ulber, 2020). Our gene ontology analyses (Supplementary S7) similarly failed to identify a dominant explanatory metabolite or pathway, suggesting that resistance emerges from multilayered interactions rather than a simple metabolic determinant. Another complicating factor lies in the nature of our feeding assay, which captures short-range interactions without being able to fully distinguish antibiosis from antixenosis (Williams & Cook, 2010). Short-range resistance may be governed by leaf-surface features, cuticular wax composition, or gustatory stimuli detectable only upon contact – traits scarcely characterized in *B. oleracea* but known to influence the behavior of flea beetles (*Phyllotreta cruciferae* L.; Bod-naryk, 1992). The absence of clear physiological cues in our dataset underscores the need for future investigation to integrate metabolomics, micromorphology, insect behavior, and transcriptomics.

### 2. Paraphyly is to be expected for polygenic architecture, especially for CSFB resistance

Identifying the genomic determinants of CSFB resistance proved challenging, but the difficulty did not stem from methodological shortcomings. In fact, throughout the paper we evaluated potential technical limitations – most notably those associated with pool-seq GWAS – and found no evidence that they undermined the analyses. The pool-seq GWAS produced stable signals, showed strong concordance with F-statistics scans, and was further supported by the high predictive ability of genomic models. Pool-sequencing itself is well established as a reliable approach for estimating allele frequencies when coverage is adequate (Schlötterer et al., 2014). Likewise, the quantitative genetic analyses were embedded in a rigorous hierarchical framework (Gautier et al. 2024), which enabled precise estimation of genic additive variance.

The real difficulty therefore arose from a genuine mismatch between phenotype and phylogeny. Resistance did not map cleanly onto the species tree: wild accessions were not uniformly resistant, nor were domesticated accessions uniformly susceptible. Instead, resistance exhibited a layered, paraphyletic distribution: part of the variation aligned with phylogenetic structure (e.g., most var. *capitata* lines being susceptible), while another component spanned taxonomic boundaries (e.g., feral populations being resistant despite their genetic proximity to var. *capitata*). Given the long-standing coevolution between *P. chrysocephala* and *Brassica*ceae – estimated at ∼16 Ma (Gikonyo et al., 2024) – it is indeed expected that different lineages repeatedly reassembled resistance through redundant pathways. Under such conditions, CSFB pressure constitutes a paraphyletic selective pressure, which can also be viewed as a fluctuating selection (Bell, 2010), considering the back and forth with feralization events. This pattern persisted even when alternative tree topologies were considered (Cai et al., 2022; Saban et al., 2023), as long as the feral origin of the “wild” populations is taken into account (Mabry et al., 2021; Cai et al., 2022). Altogether, the phenotype is best interpreted as the repeated sorting of ancestral standing variation across lineages, rather than a simple lineage-specific segregation of resistance alleles – granted that the tree structure is adequate for such a phylogeny, as opposed to a graph due to admixtures.

Such patterns are to be expected for highly polygenic traits (Goldstein & Holsinger, 1992; Yeaman, 2015; Barghi et al., 2019), where CSFB resistance fits into this framework – consistent with the broader literature on plant-insect resistance (Kliebenstein 2014; Ochoa-Lopez et al., 2018; Hervé, 2018). Indeed, when multiple loci can substitute for one another, i.e., genetic redundancy, populations can converge on similar resistance levels through many distinct allele combinations. In such systems, the same phenotypic optimum can be reached via numerous genetic paths (Höllinger et al., 2019). Under this view, the repeated “emergence” of resistant phenotypes across wild, feral, and domesticated lineages reflects independent local shifts in allele frequencies from shared standing variation rather than repeated *de novo* mutations. Paraphyly is the end point of this gradual divergence through frequency shift.

In such a context, because many loci are exchangeable in their contribution to resistance, divergent selection does not necessarily produce measurable allele-frequency differentiation: the trait can drift along multiple genetic axes without accumulating large F_ST_ values at individual loci. As a result, classical Q_ST_–F_ST_ comparisons lack power, and globally all population-genetic inference breaks down when Identity-by-State (IBS) fails to translate Identity-by-Descent (IBD). Our dominance analysis exemplifies this challenge. The major-effect QTL on C01 emerged only after incorporating a dominance component into the model, and although the overall gain in predictive ability was modest (0.688 to 0.722), the dominance-based PCoA revealed a pronounced one-dimensional – as expected (Vitezica et al., 2013) – separation between wild and domesticated accessions and likely captures what the additive Q relationship matrix could not. Specifically, dominance captured the sharing of heterozygosity excess.

## Conclusion

The paraphyletic selective pressure exerted by *P. chrysocephala* has resulted in a similarly paraphyletic segregation of resistance across *B. oleracea* lineages. Because the underlying genetic architecture is highly polygenic, neither wild nor domesticated forms occupy a stable resistance optimum. Instead, diversification and feralization acted as successive episodes of allele-frequency shift in a direction each time new and potentially independent of the previous one. This implies that feral populations are not a reservoir of ancestral resistance, but (re)attained higher resistance from a relatively susceptible background. A major-effect QTL on chromosome C01, carrying resistance alleles in feral populations and susceptible ones in var. *capitata*, appears to have undergone a recent selective sweep during this (re)acquisition of resistance, although both its physiological basis and the precise timing of the sweep remain unresolved. Our findings underscore the importance of integrating precise demographic history into quantitative genetic analyses when investigating the evolution of resistance, especially under a domestication context and a long-standing genus-specific herbivore pressure. Appreciating this history reveals the evolutionary dynamism of feral populations, previously underappreciated, yet valuable reservoir of adaptive alleles for *Brassica* breeding.

## Supplementary materials

Supplementary analyses (S1 to S7), figures (S1 to S8) and table (S1).

## Competing interests

The authors declare no conflicts of interest.

## Acknowledgments

We are grateful to the Genetic Resource Center BrACySol for providing the seeds, with special thanks to S. Théréné. We also thank the GenOuest bioinformatics platform for their technical support. This research was supported by funding from France Agrimer, SELEOPRO and the Plan Sortie Phosmet. Finally, we extend our sincere appreciation to the staff who managed the plant material, particularly L. Charlon and F. Letertre.

## Author contributions

MT, MJMD, CR, SF, AG designed the study; CF, CL, CB, DR, AL performed experiments and contributed data; MT and MB analyzed the data; MT drafted the manuscript.

## Data availability

The short-read resequencing data that support the findings are available in NCBI’s sequence read archive (SRA) under the project PRJNA1174687 (Tiret et al., 2024), and on the European Nucleotide Archive (ENA) at EMBL-EBI under accession number PRJEB105186 and PRJEB105183, and under accession numbers PRJEB91565, PRJEB91569, and PRJEB91561 for C102 (aka. PBD11), Nd125 (aka. PBD15) and Bos01 (aka. PBD20) respectively (Falentin et al., 2025). The genetic files (vcf), the raw phenotype dataset, and the scripts are freely available at: “Replication Data for: The Paraphyletic Origins of Genetic Resistance to Cabbage Stem Flea Beetle in Brassica oleracea”, https://doi.org/10.57745/4ZWIYG, Recherche Data Gouv, DRAFT VERSION, UNF:6:2vUhqizsVj/UF- b7SRM6lUA== [fileUNF]. For the BrasExplor accessions, the availability of seeds are detailed in Falentin et al. (2024).

## Notes

https://doi.org/10.57745/4ZWIYG

### Competing Interest Statement

The authors have declared no competing interest.

### Summary of Updates

Tone down conclusion and explcitly state the limid of the deployed methods.

