## Supplementary Materials for "The Paraphyletic Origins of Genetic Resistance to Cabbage Stem Flea Beetle in *Brassica oleracea*"

### S1. Bioinformatics and variant filtering

Variant calling was performed using the standard BCFtools workflow, to which we added three INFO fields beyond the default BCF fields:

- Quality-by-depth: INFO/QD = QUAL × NS / Σ(AD)
- Strand bias (Fisher’s exact test, phred-scaled): INFO/FS = phred(FS)
- Average strand position bias across pools: INFO/SP = AVG(FORMAT/SP)

Post-calling filtering was applied only on the concatenated VCF, using thresholds defined from the empirical distribution of a 100,000 SNP subset:

- Variant-level filters (SNPs and INDELs):
  - Remove variants with: QUAL < 30, MQ < 40, DP < 5, DP > 80, FS > 40, QD < 2, SP > 50, MQ0F > 0.1.
- SNP-specific filters:
  - Remove SNPs with MAX(AD) ≤ 3.
  - Remove strands/positional artefacts: |MQBZ| > 8, |RPBZ| > 4, |BQBZ| > 8, |SCBZ| > 4.
  - Remove SNPs within 5 bp of an INDEL.
- INDEL-specific filters:
  - Remove INDELs with IMF < 0.1.
  - Remove INDELs with |RPBZ + SCBZ| > 8. ◦
  - Remove INDELs within 10 bp of another INDEL.
- Population-level filters:
  - Remove variants when less than five pools show VAF < 0.1 or VAF > 0.9 (retaining variant only if there are pools with extreme frequencies), preventing leverage effects in downstream GWAS.
- Genotype-level filters:
  - Set individual genotypes to missing if GQ < 20, DP < 5, or SP > 50 (pool genotypes unaffected).

Only biallelic SNPs and INDELs with F_MISS < 0.05 were kept. For Pool-seq downstream analyses, allele frequencies were discretized into increments of 1/30, reflecting the 30 individuals per pool. The full filtering script is publicly available at: https://doi.org/10.57745/4ZWIYG

### S2. Alternative measures of population differentiation

In the main text, we relied on a Q matrix derived from the per-morphotype average of the GRM, because this representation reproduced most consistently the phylogenetic relationships previously reported for *Brassica* *oleracea* (Cai et al., 2022). In particular, it correctly resolved the clustering in *botrytis/italica* and *capitata/acephala*, and the clustering of wild accessions as feral populations. To ensure that our choice of Q did not bias inference, we compared multiple measures of population structure.

**ADMIXTURE.** Ancestry membership were inferred using ADMIXTURE v1.3.0 (Alexander et al., 2009). Runs with K = 6–10 were tested, and the minimum CV error occurred at K = 7. ADMIXTURE performed comparably to NMF in separating wild vs domesticated lineages, but was less effective at capturing divisions within var. *acephala* and wild populations, as it was applied on an approximate genotype (since ADMIXTURE does not model pool-seq sampling).

**Non-negative Matrix Factorization (NMF).** NMF was computed using the ‘NMF’ R package (v0.28; Gaujoux & Seoighe, 2010), directly on allele-frequency matrices, making it suitable for pool-seq data. Optimal rank was estimated with ‘nmfEstimateRank()’, which indicated a rank of 7 to optimize dispersion. We therefore run NMF with K = 7 and 10 independent runs. NMF was able to separate the *acephala* lineage from wild populations, but produced a highly ladderized topology of phylogenetic tree.

**Nei’s Genetic Distance.** Pairwise population differentiation was also computed using standardized Nei’s distance (Nei, 1972), adapted for pool-seq following Katada et al. (2004):

d_ij_ = -log( S_ij_/sqrt(S_ii_.S_jj_) ),

with S = (X/2)^T^(X/2) + (1-X/2)^T^(1-X/2), where X is the genotype table. Cluster-level distances were obtained by aggregating genotypes within morphotypes to produce X^S^, then recomputing d^S^. Nei-based trees resolved the wild populations close to the outgroup (*Brassica* *montana*), contrary to expectations.

### S3. Model diagnostics of the mixed models

Several bioassay protocols were preliminarily tested (not detailed here), including cotyledon assays, detached leaf assays, and exposure to cabbage stem flea beetles (CSFB) at various developmental stages. The final protocol of non choice experiment was selected because it maximized the contrast between *B. napus*, *B. oleracea* and *B. rapa*, the latter of which is known for its marked susceptibility to CSFB (Döring & Ulber, 2020; Li et al., 2024).

After scale transformation (mm^2^ → mm), phenotype skewness improved (1.672 → 0.451). Outlier removal and replication filtering resulted in:

• 742 observations, 113 genotypes.

• Mean replication 6.56 ± 1.08 (balance = 0.165).

• Fill rate = 72.9%.

### S4. Rationale for using the Q matrix in the QK model

We compared four variance-component structures:

| **Model** | **Q matrix** | **Dominance** | **Variance of ‘SSP’** |
| --- | --- | --- | --- |
| Q + K | Yes | Yes | 0.56 |
| Q + K | Yes | No | 0.00 |
| K only | No | Yes | 0.05 |
| K only | No | No | 0.02 |

Models excluding Q systematically underestimated subspecies differentiation, while dominance alone produced spurious hierarchical structures (first PC of D explained 89.4% variance). With the QK model, SSP estimates were stable. Dominance and additive effects were strongly collinear (high VIF), we then only considered the part of the dominance matrix that was orthogonal to the additive relationship matrix. Residual diagnostics showed no strong omitted-variable bias, QQ-plot consistent with Gaussian errors, small (but not null) heteroscedasticity at low fitted values, and a few influential points after filtering.

### S5. Quantitative genetics and GWAS Diagnostics

**Bootstrap vs cross-validation (CV).** Bootstrapped BLUPs and CV (80/20 resampling, n = 1000) showed a high correlation of allelic effects (ρ = 0.985), of BLUP (ρ = 0.999), and of morphotype effects (ρ = 0.925). Both converged to the same QTL peaks.

**Genomic inflation, LOCO, and local score.** We tested several GWAS methods:

- Mixed-model GWAS.
- Correction by genomic control (λ_median_; Listgarden et al., 2012).
- LOCO (leave-one-chromosome‐out) as implemented in GCTA (Yang et al., 2011).
- Local score (ξ = 3; ‘localScore’ R package v2.0.3; Simon et al., 2025).

As a result, genomic inflation correction changed no significance (ρ > 0.999), LOCO led to worse predictive ability (PA = 0.596 [0.279–0.855]), and local score was unstable across bootstrap (and high level of significance despite random permutations).

### S6. Independent validation experiment

A validation screen using *B. oleracea*, *B. rapa*, *B. napus*, and *Sinapis* *alba* confirmed strong genotype-level repeatability:

- Overall correlation with main dataset: ρ = 0.693.
- *B. napus*: 0.825.
- *S. alba*: 1.00.
- *B. oleracea*: 0.636.
- *B. rapa*: 0.599.

This experiment was used as an external validation set for GWAS signal consistency, and included the accessions ‘BO_E_LARE_W_A’, ‘BO_F_HERB_L_A’, ‘BO_F_TILL_L_A’, ‘BO_F_VALE_W_A’, ‘BO_F_BRUN_W_A’, and ‘BO_F_CHEN_W_A’.

### S7. Gene ontology enrichment

Functional annotation used the *B. oleracea* ‘HDEM’ reference genome:

- Annotation via Blast2GO v1.5.1 (Götz et al., 2008).
- GFF3 validation with ogs_check v0.4.
- GO handling with GO.db v3.18.0 (Carlson, 2023) and GSEABase v1.64.0 (Morgan et al., 2023).
- Slimming using goslim_plant OBO (2025-06-01).
- Enrichment via Fisher’s exact test.

No GO term reached the Bonferroni threshold in the significant QTL intervals (p < 8.77 × 10^-5^).

Among the top 200,000 SNPs, enriched categories (p < 0.001) included:

- Kinase activity (GO:0016301)
- Multicellular organism development (GO:0007275)
- Plasma membrane (GO:0005886)

Focusing on four distinct GO terms: ronly the response to stress go terms significantly enriched (p < 0.001).

Among targeted GO categories (response to biotic stimulus, GO:0009607; response to external stimulus, GO:0009605; response to stress, GO:0006950; secondary metabolic process, GO:0019748), only response to stress (GO:0006950) was significantly enriched (p < 0.001).

### Supplementary Figures and Tables

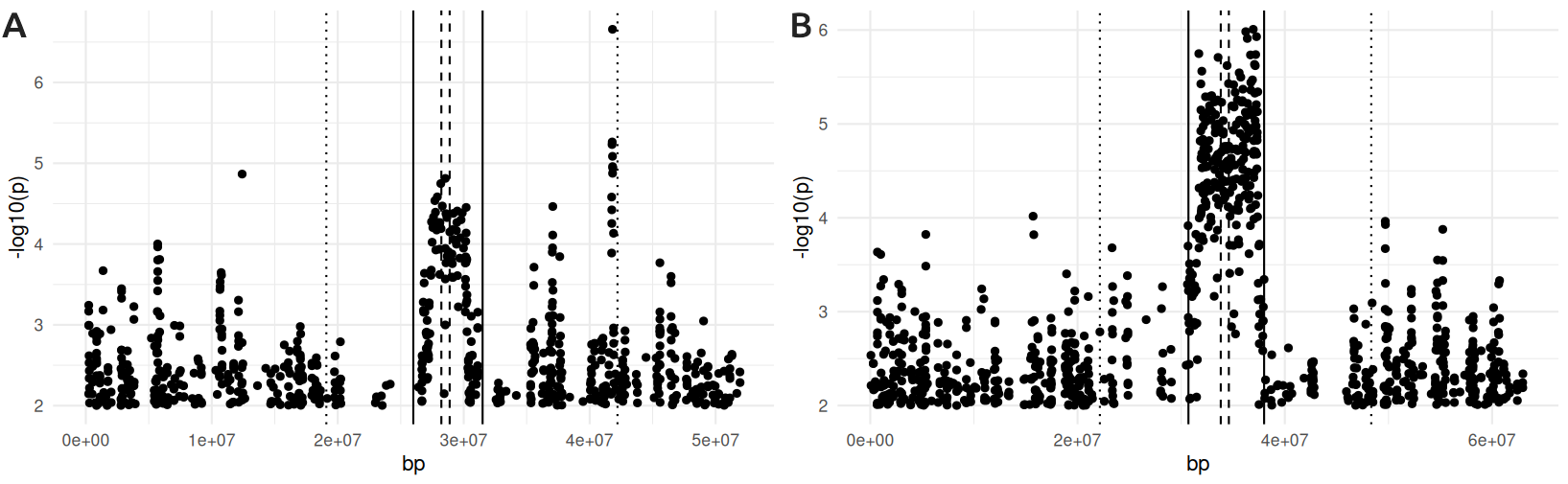

**Figure S1.** Unexpectedly high calling rate in centromeric regions of C1 (A) and C9 (B), which was visible on the Manhattan plot. Those regions were removed in the subsequent analyses.

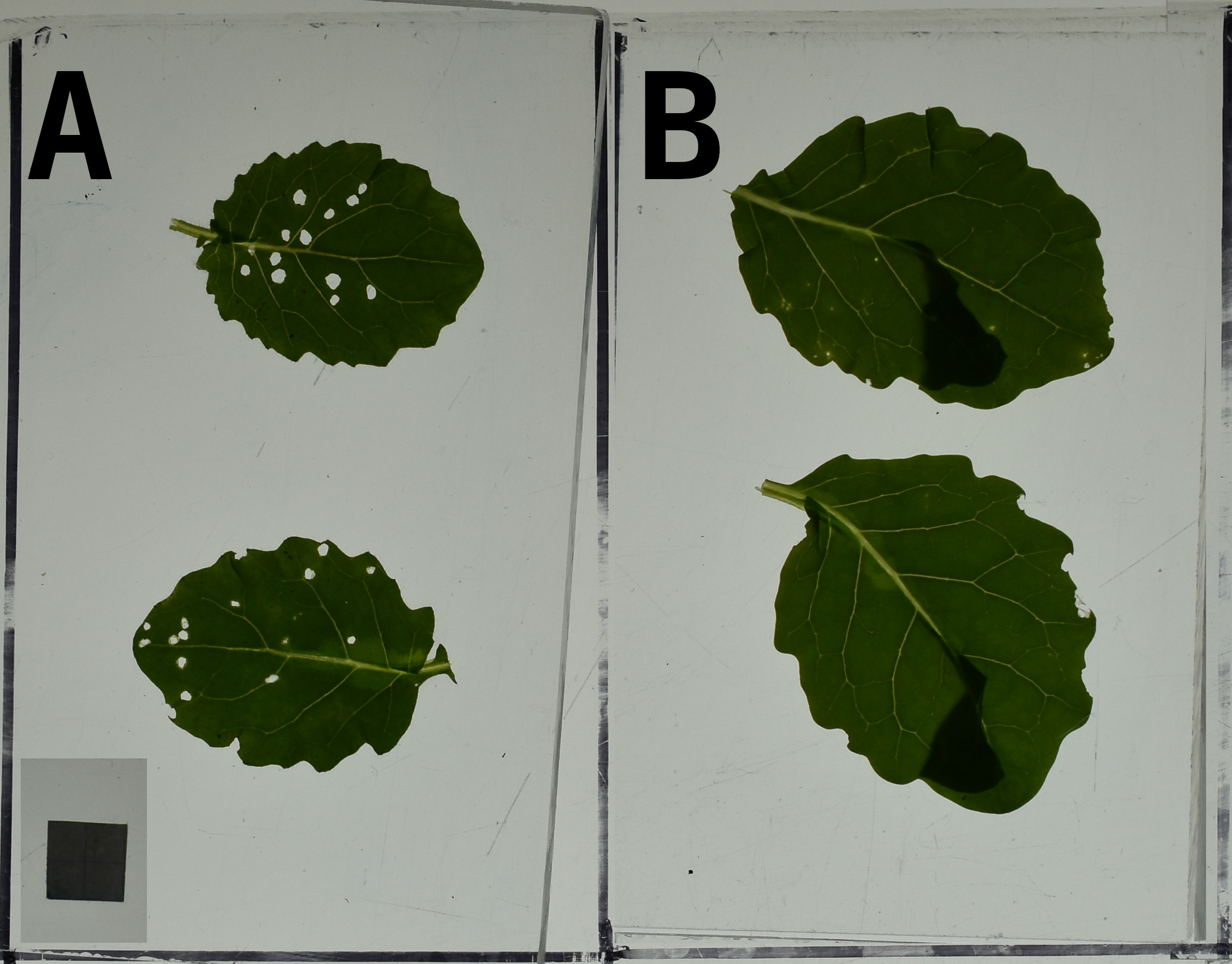

**Figure S2.** Images of different genotypes after the experiment, with the square representing 1cm^2^.(A) image of a sensitive *B. oleracea*'s leaf (var. HDEM). (B) image of a resistant *B. oleracea*'s leaf (var. BO_E_LARE_W_A).

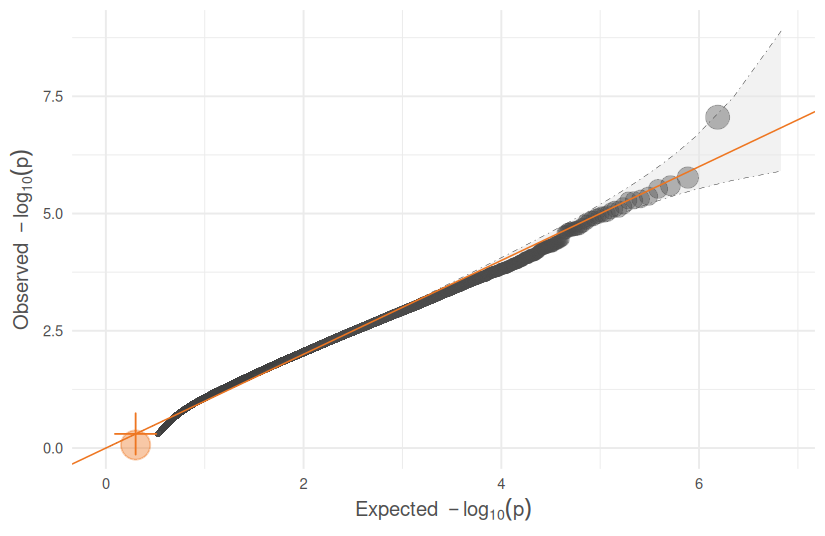

**Figure S3.** QQ plot of the GWAS.

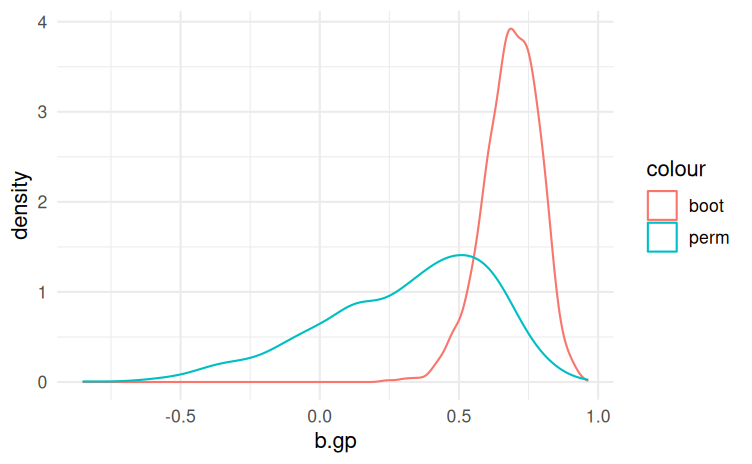

**Figure S4.** Distribution of predictive abilities across bootstrap replicates (red line) and permutation replicated (blue line).

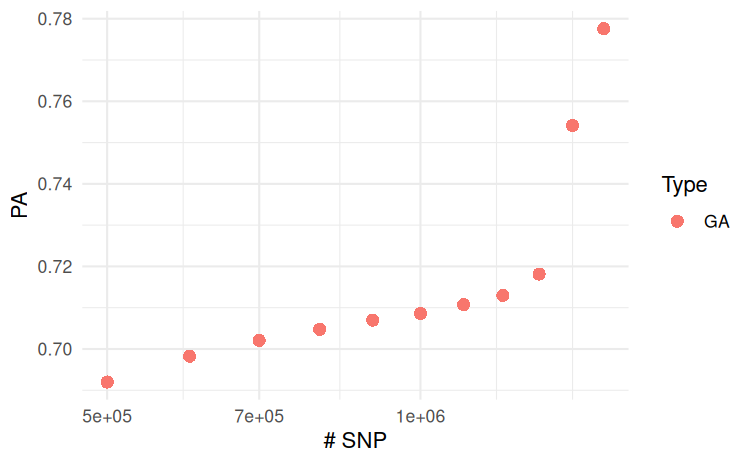

**Figure S5.** Predictive ability (PA) as a function of the SNP subset size (taking the best SNPs). The values of PA are inflated, because they are estimate from the mean allelic effect, as opposed to the core text, where the reported PA is first computed for each bootstrap replicate then averaged.

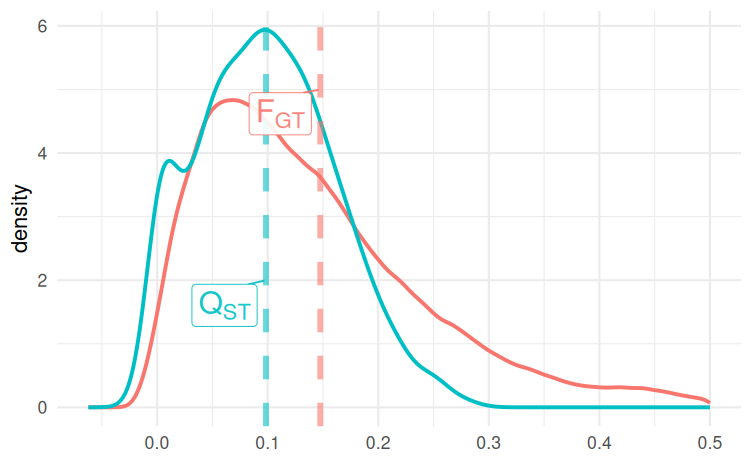

**Figure S6.** Q_ST_ vs F_GT_, phenotypic differentiation (distribution across bootstraps, blue line) and genetic differentiation (distribution across the genome, red line). Q_ST_ is higher but not significantly than F_GT_, with a plateau distribution – suggesting its instability in the estimation.

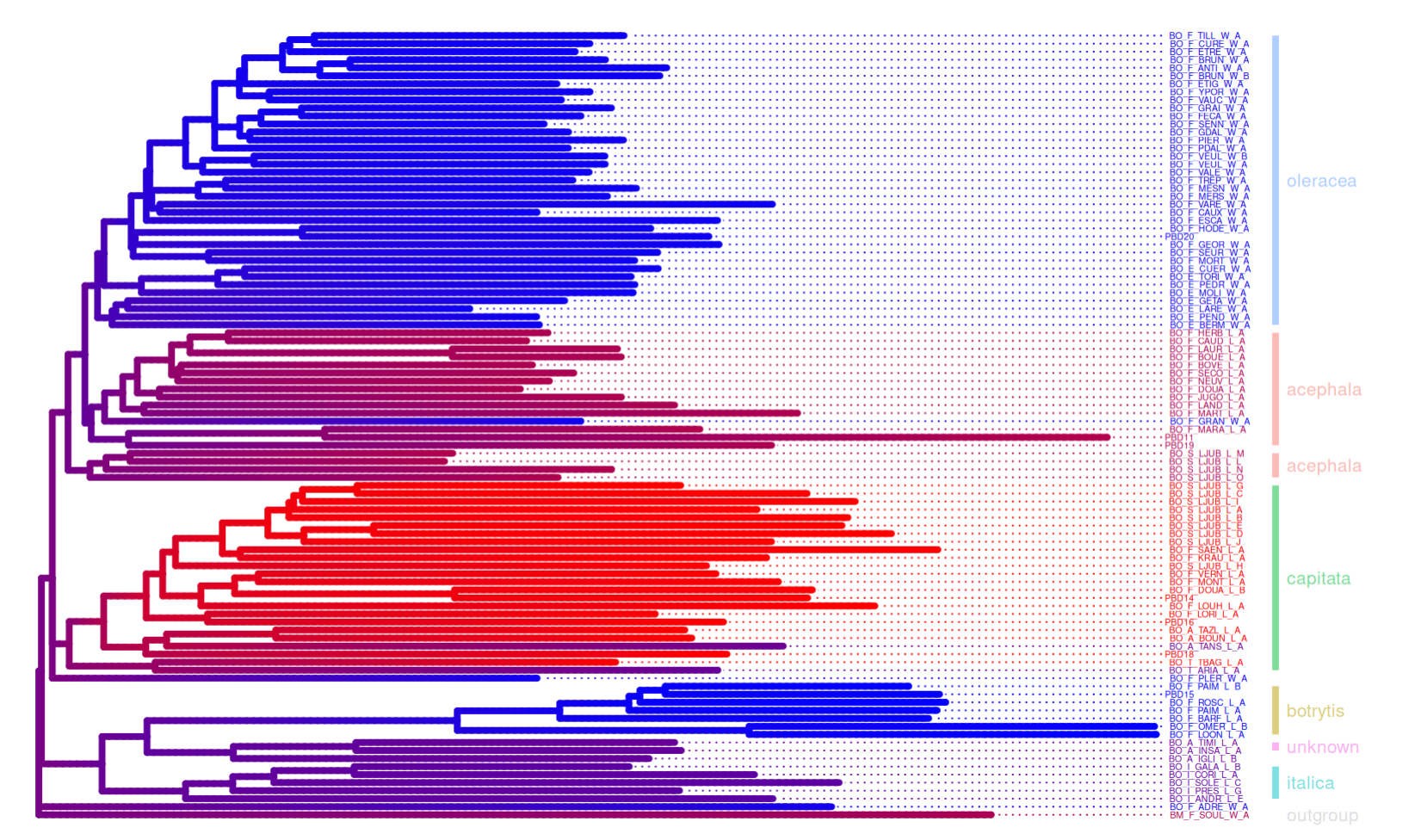

**Figure S7.** Phylogenetic tree with resistance estimated by the C1 QTL projected: resistance (blue) and sensitivity (red).

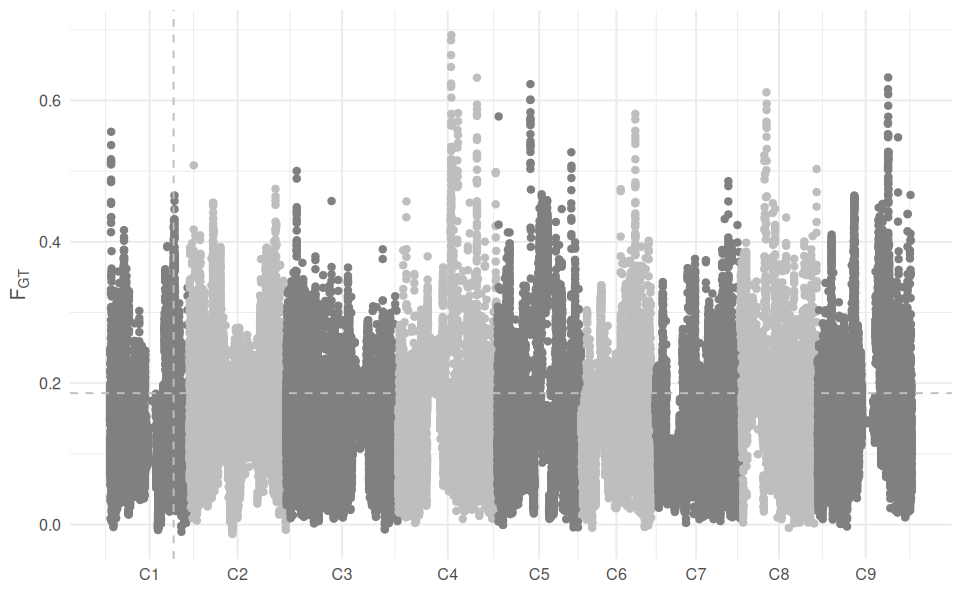

**Figure S8.** F_GT_ scan across the genome with the permutation threshold (horizontal grey dotted line) and the position of the C1 QTL (vertical grey dotted line). Most of the F_GT_ is much larger than the permutation threshold, supporting the biological significance of the morphotype clustering.

| ID | ssp |
| --- | --- |
| PBD11 | acephala |
| PBD14 | capitata |
| PBD15 | botrytis |
| PBD16 | capitata |
| PBD19 | acephala |
| PBD20 | oleracea |
| CGN18431 | capitata |
| CGN06903 | oleracea |
| CGN07077 | capitata |
| CGN07123 | capitata |
| CGN11073 | botrytis |
| CGN11132 | acephala |
| CGN15129 | botrytis |
| HRIGRU 8694 | oleracea |
| CGN18947 | oleracea |
| HRIGRU 7343 | oleracea |
| HRIGRU 8658 | italica |
| HRIGRU 6431 | acephala |
| HRIGRU 9846 | acephala |
| HRIGRU 7795 | oleracea |
| BO_A_IGLI_L_B | unknown |
| BO_A_BOUN_L_A | capitata |
| BO_A_INSA_L_A | unknown |
| BO_A_TANS_L_A | unknown |
| BO_A_TAZL_L_A | capitata |
| BO_A_TIMI_L_A | unknown |
| BO_E_GETA_W_A | oleracea |
| BO_E_LARE_W_A | oleracea |
| BO_E_MOLI_W_A | oleracea |
| BO_E_PEND_W_A | oleracea |
| BO_E_TORI_W_A | oleracea |
| BO_F_PIER_W_A | oleracea |
| BO_F_ANTI_W_A | oleracea |
| BO_F_PAIM_L_A | botrytis |
| BO_F_BARF_L_A | botrytis |
| BO_F_BOUE_L_A | acephala |
| BO_F_CAUD_L_A | acephala |
| BO_F_LAUR_L_A | acephala |
| BO_F_DOUA_L_B | capitata |
| BO_F_GRAN_W_A | oleracea |
| BO_F_JUGO_L_A | acephala |
| BO_F_KRAU_L_A | capitata |
| BO_F_LOON_L_A | botrytis |
| BO_F_MART_L_A | acephala |
| BO_F_MESN_W_A | oleracea |
| BO_F_MONT_L_A | capitata |
| BO_F_MORT_W_A | oleracea |
| BO_F_OMER_L_B | botrytis |
| BO_F_PAIM_L_B | botrytis |
| BO_F_PLER_W_A | oleracea |
| BO_F_ROSC_L_A | botrytis |
| BO_F_SEUR_W_A | oleracea |
| BO_F_TREP_W_A | oleracea |
| BO_F_VAUC_W_A | oleracea |
| BO_F_VEUL_W_A | oleracea |
| BO_F_ETIG_W_A | oleracea |
| BO_F_VALE_W_A | oleracea |
| BO_F_ETRE_W_A | oleracea |
| BO_F_ADRE_W_A | oleracea |
| BO_F_BRUN_W_A | oleracea |
| BO_F_BRUN_W_B | oleracea |
| BO_F_CAUX_W_A | oleracea |
| BO_F_CURE_W_A | oleracea |
| BO_F_ESCA_W_A | oleracea |
| BO_F_FECA_W_A | oleracea |
| BO_F_YPOR_W_A | oleracea |
| BO_F_GDAL_W_A | oleracea |
| BO_F_HODE_W_A | oleracea |
| BO_F_MERS_W_A | oleracea |
| BO_F_PDAL_W_A | oleracea |
| BO_F_SENN_W_A | oleracea |
| BO_F_TILL_W_A | oleracea |
| BO_F_VARE_W_A | oleracea |
| BO_F_VEUL_W_B | oleracea |
| BO_S_LJUB_L_B | capitata |
| BO_F_GEOR_W_A | oleracea |
| BO_I_CORI_L_A | italica |
| BO_I_ANDR_L_E | italica |
| BO_I_PRES_L_G | italica |
| BO_I_SOLE_L_C | italica |
| BO_S_LJUB_L_A | capitata |
| BO_S_LJUB_L_D | capitata |
| BO_S_LJUB_L_C | capitata |
| BO_S_LJUB_L_O | acephala |
| BO_S_LJUB_L_E | capitata |
| BO_S_LJUB_L_G | capitata |
| BO_S_LJUB_L_H | capitata |
| BO_S_LJUB_L_J | capitata |
| BO_S_LJUB_L_L | acephala |
| BO_S_LJUB_L_M | acephala |
| BO_S_LJUB_L_N | acephala |
| BO_T_ARIA_L_A | unknown |
| BO_T_TBAG_L_A | capitata |
| BO_F_HERB_L_A | acephala |
| BO_F_NEUV_L_A | acephala |
| HDEM | italica |
| BO_E_PEDR_W_A | oleracea |
| BO_E_BERM_W_A | oleracea |
| BO_F_GRAI_W_A | oleracea |
| BO_F_SAEN_L_A | capitata |
| BO_F_DOUA_L_A | acephala |
| BO_F_LAND_L_A | acephala |
| BO_F_LORI_L_A | capitata |
| BO_F_LOUH_L_A | capitata |
| BO_F_VERN_L_A | capitata |
| BO_F_SECO_L_A | acephala |
| BO_I_GALA_L_B | italica |
| BO_S_LJUB_L_I | capitata |
| CGN14063 | capitata |
| BO_F_BOVE_L_A | acephala |
| PBD18 | capitata |
| BO_E_CUER_W_A | oleracea |
| BO_F_MARA_L_A | acephala |

**Supplementary Table S1**: list of accessions and subspecies.
